# Learning massive interpretable gene regulatory networks of the human brain by merging Bayesian Networks

**DOI:** 10.1101/2020.02.05.935007

**Authors:** Nikolas Bernaola, Mario Michiels, Pedro Larrañaga, Concha Bielza

**Affiliations:** Computational Intelligence Group, Departamento de Inteligencia Artificial, Universidad Politécnica de Madrid, Madrid, Spain

## Abstract

We present the Fast Greedy Equivalence Search (FGES)-Merge, a new method for learning the structure of gene regulatory networks via merging locally learned Bayesian networks, based on the fast greedy equivalent search algorithm. The method is competitive with the state of the art in terms of the Matthews correlation coefficient, which takes into account both precision and recall, while also improving upon it in terms of speed, scaling up to tens of thousands of variables and being able to use empirical knowledge about the topological structure of gene regulatory networks. We apply this method to learning the gene regulatory network for the full human genome using data from samples of different brain structures (from the Allen Human Brain Atlas). Furthermore, this Bayesian network model should predict interactions between genes in a way that is clear to experts, following the current trends in explainable artificial intelligence. To achieve this, we also present a new open-access visualization tool that facilitates the exploration of massive networks and can aid in finding nodes of interest for experimental tests.

## Introduction

With the advent of high-throughput measurement technologies in biology in the 1990s, such as in situ hybridization [1] [2] or RNA microarrays [3], it has been possible to collect information for tens of thousands of genes from every tissue sample or even at the level of a single cell [4]. Since most of the information about the development and function of every living being is codified in its genome, we can study the level of expression of each gene in different conditions. This makes it possible to reconstruct the underlying regulatory relationships between the genes and, therefore, get closer to understanding their function [5] [6]. Due to the combinatorial nature of gene regulation [7] and the size of the genome (which can have tens of thousands of genes), it would be intractable to experimentally determine all of the regulatory links. Even further, a complete model would have to take into account post-transcriptional modifications. To solve this problem, many computational methods have been proposed to infer the gene regulatory network (GRN) from expression data [8] [9]. The models learned can then be used to guide biological research by letting researchers test the interactions predicted by the network. The main objective of this paper is to present an algorithm capable of reconstructing the GRN for the whole human genome using gene expression data from the human brain to obtain a model that is accurate, easily interpretable and capable of quantitative prediction of the levels of gene expression. To accomplish this, we first surveyed some of the most common methods for learning GRNs and listed their main advantages and limitations. We decided to use Bayesian networks (BNs) (BNs) [10] [11] due to their superior interpretability and then subsequently present an algorithm, FGES-Merge, based on the fast greedy equivalence search (FGES) method [12] that solves the usual problems that BNs have when dealing with very large networks. Our contribution is adapting the model to cases with many more variables and to the concrete topology of GRNs were some nodes are very densely connected and violate the assumption of sparsity in previous algorithms.

To facilitate the use of our model as a research aid, we also present a new open-access visualization tool, NeuroSuites-BNs, that can easily represent networks with tens of thousands of genes and allows the user to focus on nodes of interest and graphically perform the usual operations on BNs (introducing evidence, making probabilistic inference, showing the parents, children and Markov blanket of any variable, etc.).

In the next section, we introduce the basic biological knowledge needed to formulate the problem of reverse engineering a GRN and we will survey some of the most common methods to solve it. We then emphasize the work done with BNs along with their main advantages and limitations. Afterwards, we formulate the problem of learning and visualizing a genome-wide GRN for the human brain and present our method for solving it. Then, we show the results of applying our method to the DREAM5 GRN benchmark and the Allen Human Brain Atlas dataset, including the visualization of the reconstructed networks using FGES-Merge and Neurosuites-BNs. Finally, we present our conclusions and possible improvements to the work.

## Literature review

In this section, we will introduce the biological background to the problem of reconstructing GRNs from data. Then, we will briefly summarize some of the most important methods for learning GRNs. Finally, we will introduce BNs as the model of choice for the problem at hand. discuss their theoretical background, previous work done with BNs in the field and the main limitations we need to overcome to solve the problem of inferring a full genome network from data for the human brain.

### Genetics

One of the most important scientific advancements of the last century was the discovery that all heritable information in a living organism is encoded biochemically and stored in chromosomes, very long polymers of double stranded, helical DNA [13]. Information stored in DNA can be dynamically read in a process that is also biochemically consistent among most species. One of the DNA strands is transcribed into RNA, a single stranded polymer of nucleic acids, which acts as an information carrier that is in turn translated into proteins in ribosomes. Proteins are polymers of amino-acids that, depending on the folding structure, can carry out almost all cellular functions. The flow of information through the cell (see Fig. 1) is of the uttermost importance in biology and has been named the *central dogma of molecular biology* [14]. One of the most important facts about gene expression is that every RNA strand encodes one, and only one, protein. This is because every triplet of nucleic acids in the RNA strand maps to a unique amino-acid in a way that is consistent across species. This mapping is called the *genetic code* [15].

**Fig 1.**
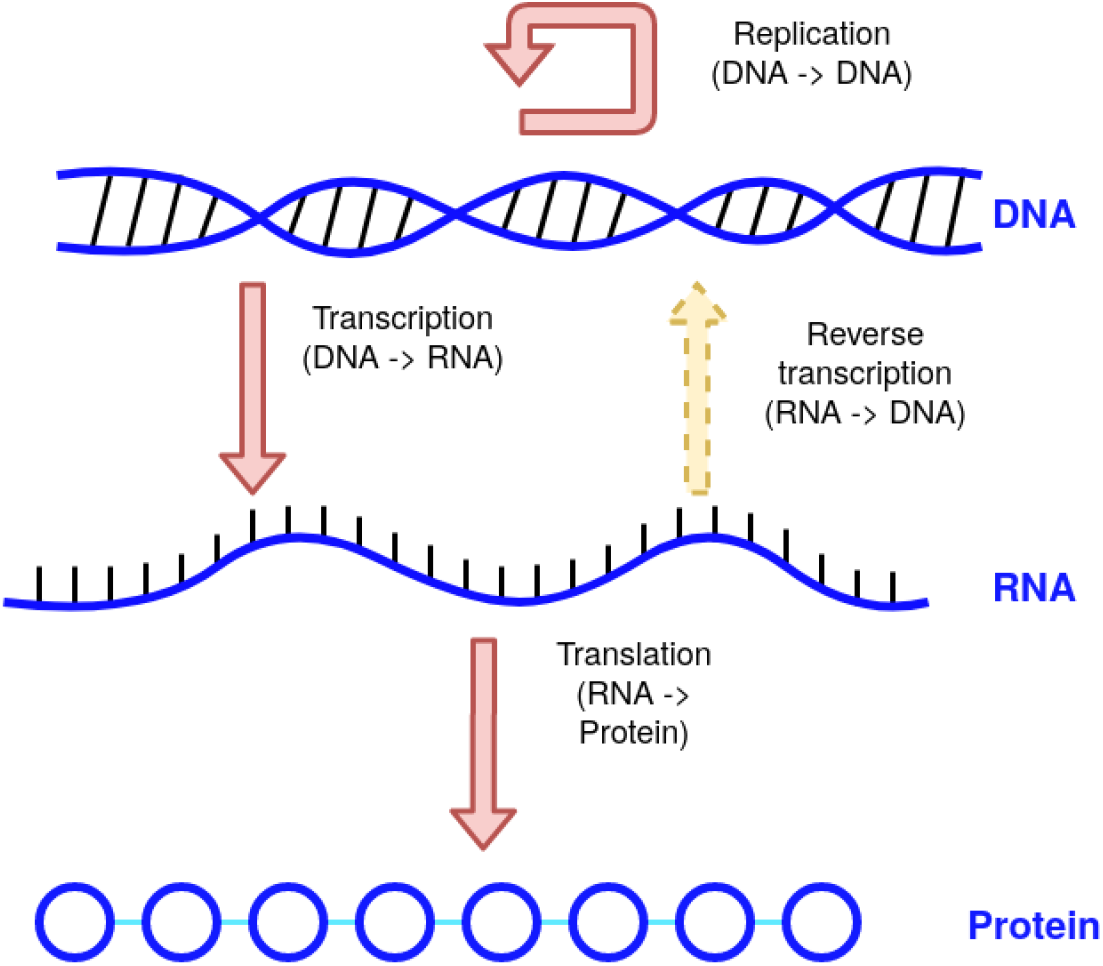
Flow of information in a cell. Diagram explaining the central dogma of molecular biology. (Reverse transcription is very error prone and only seen in some virus species.)

The full complexity of every living organism has to arise from the interactions between these biochemical components. Therefore, there is a strong scientific interest in understanding these interactions both to improve our fundamental understanding of how life arises from biochemistry and because of the important practical applications in medicine and drug manufacturing. The main problem is that probing live cells to measure interactions directly is incredibly difficult, so studying them directly is almost out of the question. However, thanks to the genetic code, we can substitute the questions about protein interactions with questions about the concentration of the messenger RNA (mRNA) that encodes for each of the proteins. In addition, thanks to advances in high-throughput sequencing technology during the last two decades, measuring the abundances of different biochemical components (including mRNA) is much more manageable, making it possible to do so at a very large scale for relatively low cost. The abundances do not give us the full picture of the interactions, but this trend in the availability of data has provided a powerful motivation to try to computationally reconstruct the interaction structure that underlies gene expression. These structures are called GRNs, and their reconstruction is one of the central efforts in the field of systems biology.

#### Data collection

As mentioned in the previous section, direct measurements of interactions at the cellular level are almost impossible and direct measurement of the protein products requires a very complex analysis pipeline which makes it less efficient than using transcriptomic measurements [16]. Microarrays and *in situ* hybridization (ISH) are the most common methods used to collect data. Microarrays are an older method that works by lining a chip with microprobes that puncture a biological sample and take a sample of the cytoplasm. Each probe in the array is lined with the complementary chain to the RNA we want to detect (a different one in each probe) in such a way that after washing out the array, only the bound RNA will be found in each probe. The bonding strands are made to induce a fluorescent molecule to emit light when bound so that each probe will emit light corresponding to the amount of binding RNA found in the sample. Via calibration, the level of light measured at each point on the chip can be mapped to a concentration level for the bound gene. The main limitations of microarrays are that we can only measure known transcripts (since the complementary strand has to be designed onto the chip) and that the microarrays give a measurement of the population of cells in the tissue sample. Since there might be many different cell types in the population, the measurement may not be representative of any single one of them. ISH is a newer method that consists of taking a sample of cytoplasm from a single cell, filtering to keep only mRNA and reverse transcribing the mRNA into DNA. Finally, PCR is used to amplify the DNA in the sample up to a level where it can be more precisely measured. By knowing the value of the amplification factor, we can then estimate the original level of RNA in the cell. Therefore, microarrays give an average expression level for the tissue sample, while ISH can give accurate, single-cell measurements [17].

This paper uses the Allen Human Brain Atlas dataset [18], which was mainly created using microarrays. Although there were also some ISH data available, they was not complete, and we decided not to include them to avoid the problems inherent in mixing two data sources (especially since we did not have ISH data for all areas of the brain).

#### Genetic regulation

The processes that regulate which genes are expressed and how much of the protein is synthesized are the topic of study of genetic regulation. In general, we are interested in seeing how different environmental conditions (internal or external) change the amount by which each protein is synthesized. Gene expression can change due to hormonal responses, any physical or chemical changes in the external environment, internal chemical changes, or the expression of other genes. The whole process of gene regulation is thus combinatorial, taking into account many factors for each gene. The process is dynamic, since it is always responding to environmental changes and feedback (usually negative), and stochastic, since even under the same conditions, we can only say that the level of expression will also be similar due to the imperfections of biological processes and the inherent instability of chemical processes.

Although the most correct model we have for the behaviour of gene expression is a system of stochastic partial differential equations [19], this model is generally intractable for the study of even the simplest of organisms if we are interested in more than a few tens of genes. One common way of simplifying the model into one that is usually good enough is to assume that all regulations can be modelled as genetic interactions. This means that we assume that any non-genetic factor (hormones, temperature, chemical changes in the environment, etc.) will not have an effect on the level of expression of genes except indirectly, via mediation by another gene. In this way, we can eliminate all external factors from our model and be left with only interactions between genes. This network structure of interactions is the GRN. In this work, we will make the additional, very common assumption [6] of taking the steady-state expression level which means we will work with static instead of dynamic models.

### GRNs and their properties

We are interested in both the topology of the network to see the interactions between genes and in accurately predicting changes in the level of expression of some genes given other changes in the level of other genes. In this section, we will briefly discuss the mathematical formalisms required to properly define these GRNs, some of the most important biological properties that can be used to constrain the space of possible structures and the multiple available methods for learning them.

#### Notation

For the representation of a GRN, we will use a directed or undirected graph 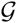 (depending on the method). 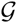 is a pair (*V, E*), where *V* is the finite set of vertices or nodes indexed by 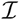 and *E* is a subset of 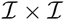, with element (*i, j*) indicating an edge between nodes *i* and *j*. If the network is undirected, then the set of edges is symmetric under swapping the indices of its members, that is, (*i, j*) ∈ *E* ⟺ (*j, i*) ∈ *E*.

In the context of GRNs, the set of nodes is always associated one-to-one with the set of genes. Edges are interpreted as relationships between genes, but the precise definition of the relationship will depend on the mathematical model being used. Nevertheless, the topological structure of the network is useful by itself, as it gives a visual intuition of the interactions at play. Some facts we can know just from the structure are the presence or absence of hubs, which are nodes with a higher than average number of edges attached, and the density of the network, which is the ratio of edges per node. We can also calibrate the structure with known relationships by checking if correct edges exist between the correct nodes.

Most models will add more information to the network, both to the nodes (usually a base level of expression, but sometimes more parameters, e.g., the standard deviation in a Gaussian BN; see below) and to the edges (e.g., regression coefficients and pairwise correlations). This information will be used to predict the changes in the level of expression along the network and to estimate the strength of relationships between genes.

We will be learning the GRNs from a dataset 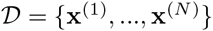 where 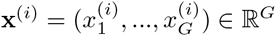, *i* = 1, …, *N*, with *N* the number of measurements and *G* the number of genes.

#### Topological properties of GRNs

Some of the most important information we can extract from the topology of the network is the degree distribution, which is given by the number of edges attached to each node of the network. In the case of directed networks, we can further distinguish between the in-degree and out-degree, which relate to the number of edges going into or out of each node, respectively. This information can be either compared to the known properties of real GRNs to see if the methods are working correctly or used beforehand to restrict the space of structures that will be searched over.

For a more in-depth overview of the topological properties of GRNs see [7], [20] and [21]. We will summarize these properties here:

- GRNs are locally dense but globally sparse: GRNs have a small number of edges compared with a fully connected network. The maximum number of edges would be G 2 (if we allow for self-edges). The actual number of edges in a GRN varies depending on the network but is typically O(G) with a small (10) constant. This means that the network is sparse. However, the degree distribution of the GRN is very fat tailed such that instead of finding that most nodes have one edge, we find many with no edges and hubs with many edges. This means that the connected components of the network are dense, but there are many disconnected components that make the network sparse. This translates to a limited number of edges in the network, and although it seems like it should make the problem more tractable, it is very problematic for many learning methods that require sparsity since, to the best of our knowledge, they usually require local sparsity.
- The in-degree distribution is a Laplace distribution with an upper limit: The in-degree of a node in a GRN is the number of regulators a gene has. This number is usually small since most genes require just one regulatory factor, although it can be higher in more complex processes that need multiple conditions to be met at the same time. However, there is a physical restriction on the number of regulators. In the case of transcription factors, the only way they can affect the expression of a gene is to be physically close to that gene, affecting the DNA directly. Since there is limited space around each gene, there can only be a limited number of transcription factors and thus a limited number of regulators. This number is lower in prokaryotes than in eukaryotes, since there are other mechanisms in eukaryotes that can be affected from a distance (e.g., histone coiling). This translates to an upper bound to the number of parents for each node.
- The out-degree distribution is scale-free distributed at the upper tail: Since the degree distribution is fat-tailed and the in-degree distribution is bounded, the out-degree distribution must also be fat-tailed. The nodes at the tail of the out-degree distribution are hubs of the network, and the biological interpretation is that they are regulators of transcription (transcription factors) used in many processes at the same time. Biologically, this is due to hubs being more evolutionarily stable. Any useful adaptation can be constructed on top of already existing regulatory machinery, which is easier than having a new regulatory network emerge simultaneously with the new adaptation.

### Methods for learning GRNs

Now that we have discussed some basics of the structure and notation used in the field of learning GRNs we will dedicate this section to summarizing the different existing approaches for the task of reconstruction with a brief explanation of some of the most important methods and current work. For a more in-depth review see [6]. We will also emphasize the advantages and disadvantages of each method and explain why, in the end, we decided to work with BNs. We will not address some more complex methods that involve combinations of multiple approaches or methods for modelling dynamic GRNs. A recent review of these methods can be found in [22].

#### Basic statistical methods

This first group of methods is based on basic statistical methods that can be used to measure dependencies between variables. They start from a fully connected network and then associate a weight to each edge. The output can then be thresholded to obtain a reasonable approximation of the topology of the network. The main advantage of these methods is that they use very common statistical techniques and so are readily available and computationally cheap. They are also reasonably accurate in finding the network topology which makes them some of the most popular.

##### Correlation networks

These methods are based on calculating the pairwise Pearson correlation (although other measures are possible) between all pairs of genes. For two random variables, *X* and *Y*, the pairwise Pearson correlation is given by their covariance normalized by the product of their variances:

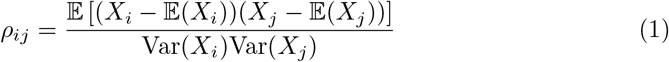

We can estimate the pairwise correlation between the columns of our dataset 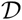 to obtain a correlation matrix, ***C*** ∈ ℝ^*G×G*^. This matrix is used to assign a weight to each of the edges of the network. Then, we can apply a threshold to the absolute value of the correlation that will depend on the level of sparsity we want to obtain for the GRN structure. Since the correlation is symmetric, the network will be undirected.

The advantages of this method are that the complexity is only linear in the number of instances and quadratic in the number of genes which makes it very popular for genome-wide or other massive studies. The underlying biological assumption is that genes with interacting regulation should have correlated expression. This is plausible, and correlation methods are consistently reliable [23]. The main limitations of correlation networks are that they fail to distinguish between direct and indirect regulation, they do not capture nonlinear interactions well, they are undirected and they are not predictive. Thus, they cannot distinguish between direct and indirect regulation since if gene *A* regulates gene *B*, which in turn regulates gene *C*, it is very likely that the correlation between *A* and *C* will be high, which would be detected as *A* directly interacting with *C*. This is worsened by the presence of hubs in GRNs since all children of a single hub (in the true GRN) will be correlated to each other and to the hub, making it truly difficult to discern which of them is the true regulator. Furthermore, the lack of direction makes it hard in general to distinguish the regulator from the regulated genes. In addition, using the Pearson correlation, which is linear, can fail to capture more complex types of interaction (even though this is not usually a problem in practice). Finally, correlation networks are not predictive since correlation is just a statistical measure of association and therefore cannot be used to make quantitative predictions about expression levels.

##### Mutual information networks

As a way to relax the assumption of linearity implicit in correlation networks some methods apply an alternative measure based on information theory. They use mutual information (MI) in the same way correlation was used in the previous method. Let *X_i_* and *X_j_* be two discrete random variables and let *P* (*X_i_, X_j_*) be their joint probability distribution. The MI between the two random variables is defined as:

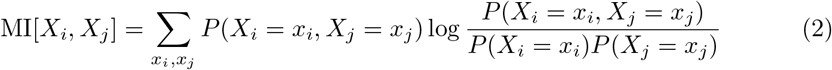

where *x_i_* and *x_j_* are the values that *X_i_* and *X_j_* can take, respectively, and *P* (*X_i_*) and *P* (*X_j_*) are the marginal probability distributions of *X_i_* and *X_j_*. MI[*X_i_, X_j_*] is zero when *X_i_* and *X_j_* are independent.

The main idea behind MI-based methods is to estimate the probability distributions from the data and then calculate the MI for each pair of genes. The resulting MI matrix gives a score to each edge in the fully connected network which will be undirected. This can be thresholded to obtain a so called relevance network [24]. Some of the most popular methods are ARACNE (the most basic) [25], CLR [26], which adds a step to ARACNE to try to reject spurious correlations and indirect influences, and MRNET [27], which takes a maximum relevance, minimum redundancy approach.

The advantage of MI networks is that they are almost as computationally cheap as correlation networks while being able to capture nonlinear relationships. The main drawbacks are that, again, they do not offer a predictive framework, that the estimation of the probability distributions will be highly sensitive to noise in the samples when the sample size is small and that they overestimate the relationships since they cannot distinguish between direct or indirect regulation.

##### Regression networks

The previous methods use basic statistical measures to compute the dependencies between genes. A different way to approach the problem is to try to predict the expression level of one gene given the remaining genes. One obvious way to do this is with a regression model, the simplest of which would be a linear regression model. In the context of GRNs, the model would be learned by regressing each gene in turn against all others and the coefficient for each gene in the regression would be used as the weight for the edges of the network. That is, each gene’s expression level *X_i_* is given by:

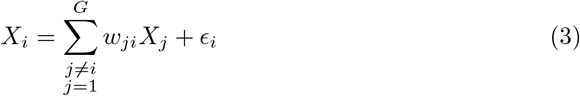

where *ϵ_i_* is a noise term. Solving the regression problem would give the weight *w_ji_* associated with the edge from gene *X_j_* to *X_i_*. Note that unlike previous methods, regression-based networks are directed (and can even have bidirectional edges). As in the other methods, a threshold can be applied to the weights of the edges to prune the network.

Regression-based methods are generally very powerful since they provide both a structure for the network and a model that can predict gene expression. They were considered to be the state of the art in the last DREAM (Dialogue for Reverse Engineering of Models) challenge [28]. In particular, TIGRESS [29] uses a linear regression method with L1 regularization to force some of the *w_ji_* to be zero which avoids using an arbitrary threshold. Another variation on this theme is GENIE3, which was also used in the DREAM5 challenge, and uses random forest regression to make the method more flexible and non-parametric. This method was first presented in [30], and then improved in [31] and [32].

Regression models are only slightly more computationally intensive than the previous methods and can, theoretically, recognize indirect dependencies between genes and assign lower coefficients to them. In practice, however, regression models tend to fail with limited data, since the highly correlated real structure of gene expression causes regression networks (even when regularized) to give spurious results in the same way as correlation and MI networks.

#### Deep learning methods

Currently, we can also find deep learning methods that outperform linear regression-based methods for inferring gene expression [33]. However, there are two downsides to these approaches: they require a large amount of data, which is not always easily available, and they do not give the structure of the network, merely a black box approach to the inference problem. However, even if they outperform the other methods, since we are interested in the structure of the network and do not have an abundance of data for the problems we wish to work on, we do not use deep learning in this study.

#### Probabilistic models

The models above work by defining some measure of dependency in a pairwise or all-to-one way. However, none of these methods explicitly defines a probabilistic model of the data (although there is an implicit model in the regression). In this section, we briefly introduce Gaussian graphical models and then review the work done with BNs, which will be the main focus of the article afterwards.

##### Gaussian graphical models

One of the simplest probabilistic models we can consider is a multivariate normal distribution. The probability density distribution for a multivariate normal vector **X** ∈ ℝ*^G^* is given by:

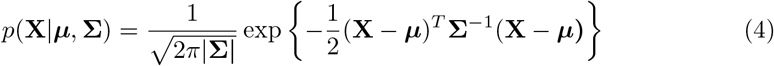

where ***μ*** is the mean vector and **Σ** is the covariance matrix.

Gaussian models give us the whole power of probabilistic inference, which allows us not only to make predictions about gene expression but also to quantify our uncertainty. Furthermore, they have a very important property in that the inverse of the covariance matrix, the precision matrix ***W*** = **Σ**^*−*1^, contains the *partial correlations* between the pairwise components of **X**= (*X*_1_, …, *X_G_*).The partial correlation is the residual correlation between two variables once the effect of all other variables has been subtracted. Therefore, it is a better measurement of the strength of the relationship between genes and is less vulnerable to making spurious associations due to the highly correlated nature of the GRN.

This result is used by Gaussian graphical models [34], which are learned by treating the measurements of expression as a multivariate normal random vector and estimating the precision matrix from the samples using maximum likelihood estimation. The number of parameters is of the order *G*^2^, so regularization techniques are used, mainly sparse regularization techniques such as L1 because they have the benefit of having a topological interpretation for building the network structure, that is, the nonzero entries of the precision matrix correspond to the edges of the underlying GRN.

Gaussian models have all the properties we want: they are probabilistic, interpretable and have both a topological and a predictive component. However, they are limited mainly by the fact that it is generally very difficult to estimate a high-dimensional precision matrix from limited data. It might even be theoretically impossible when the number of samples is less than the number of dimensions of the matrix (although an estimate can be found but with no guarantee of correctness). There is also the assumption of normality for the expression data, which implies linearity in the relationships. Even though this is a strong assumption, it is used in linear regression networks (implicitly) and Gaussian graphical models, and it seems to be a reasonable approximation that is very accurate in practice and enormously simplifies computations.

### Bayesian networks

Every method we have discussed so far uses a top-down approach to the task of learning the topology of the network, in the sense that models start with all of the variables at the same time and then prune them. The class of methods we will focus on, BNs, have a similar objective to Gaussian models in that they try to build a joint probabilistic model, but they approach it in a bottom-up way, building the model from local conditional parts.

BNs [10] are probabilistic graphical models that combine probability and graph theory to efficiently represent the probability distribution of a group of variables *G* = {*X*_1_, …, *X_n_*}. BNs model probabilistic conditional dependencies and independencies between the variables in *G* in terms of a directed acyclic graph (DAG) and a series of conditional probability distributions (CPDs) [35]. Each of the nodes in the graph represents a variable in *G* with the edges representing conditional dependencies between the variables. Each of the CPDs is associated with a variable *X_i_* and gives the probability distribution of that variable conditioned on its parents in the graph, that is, the nodes that have edges directed towards *X_i_*, which we denote **Pa**(*X_i_*). The formula for the joint probability distribution of the variables given all the CPDs is:

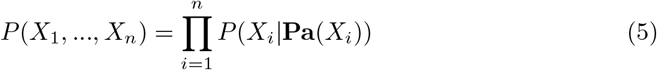

#### Previous work using BNs

BNs and other probabilistic graphical models are commonly used for other problems regarding brain data, specially fMRI, as in [36] or [37]. In the field of GRNs, one of the earliest approaches was [11] which used a simple approach of searching the whole space of structures for the one with the maximum likelihood. [38] used the PC algorithm on microarray data to obtain a GRN, and in [39], co-regulated modules of genes were first identified and then a BN was built for each of them. Some more recent advances include [40] which used a variant of the Chow-Liu algorithm to be able to learn a BN in quadratic time but with a severe limitation on the structure since it must be a polytree. Works such as [41] attempted to simplify the problem of learning the structure by including expert knowledge in the form of a prior for the structure. In [28] we can find several BN-based methods, of which the best used simulated annealing to add a stochastic element when learning the DAG structure and average the results to increase their resiliency against possible errors. In [21], the authors used topological information to restrict the space of structures and accelerate the search. Works such as [42] present a parallelized method to learn a genome-wide network that is implemented in a supercomputer. One interesting approach is [43], in which the authors learned small local networks around each node and then combined them. We also have the very recent [44], which reviews different ways of introducing prior knowledge as structural restrictions and shows that these restrictions lead to improved results on the reconstruction of the networks.

#### Advantages and limitations of BNs

BNs did not show a good overall performance in the DREAM5 challenge [28], but their interpretability makes them a good model for improvement. They readily encode the regulatory network in their graph structure, and the way they can be built avoids the problem of finding a fully connected network, as in regression or pairwise methods. This reduces the need to use an arbitrary threshold to remove some edges, since they are taken out in a less arbitrary way by testing for conditional independence, and they can capture indirect regulation well. Their probabilistic nature is especially important in this domain because it allows us to run inference through the learned structure, which is done by setting some genes as fixed evidence and then querying other genes to see how they have changed, thus providing a view on how some genes influence the expression of the rest. Furthermore, the output at each node is a probability distribution, which is a very natural way of presenting the variability inherent to the stochastic process of gene regulation.

As seen above, the main disadvantage of using BNs is that for them to be scalable to genome-wide datasets, we require either restrictions on the structure or high-performance computing (HPC). Even when restricting the structure using methods such as sparsity penalties, priors or structural restrictions, we still have to deal with the sheer size of the network we are trying to build, resulting in most of the common methods being just too slow as far as the GRN of the human brain is concerned. We also have the problem that this organ is not as well studied in other organisms, so introducing the small amount of reliable expert knowledge we have does not reduce the size of the problem much while also introducing a slowdown of having to make the algorithm consistent with this knowledge. The fastest method that does not use HPC or restrictions for gene expression data is [43], but its largest network is orders of magnitude smaller than what we need, that is, it works with networks of less than 1000 nodes while we need to build a 20.000 node network. On the other hand, methods that claim to scale to millions of variables such as FGES [12] are not applicable to our problem since they require local sparsity, which, as we saw in the section on topological properties of GRNs, we do not have.

We took the best BN method in the DREAM5 challenge and the model in [43] as the state of the art for learning GRNs of 1000+ nodes. The first used a simulated annealing algorithm with the catnet R package [45] to learn the network multiple times and merged the results to increase robustness. The second used a local approach (at the node level) to learning and then combining the resulting networks. By combining both methods with some important changes to achieve competitive times for much larger networks we arrived at our new proposal, FGES-Merge, which we present in the next section.

Finally, we want our network to offer a predictive framework for gene expression and be easy to interpret for biological research. This is of the uttermost importance these days since law changes like the General Data Protection Regulation (GDPR) and the right to an explanation from algorithmic decisions [46] make it indispensable that models that might have applications in the biological sciences and medicine are easily understood by experts. This trend towards interpretable and explainable AI is one of the main reasons we decided to use BNs in the first place and motivated the creation of our visualization tool.

## Problem statement and methods

### Problem statement

#### Learning the genome-wide GRN for the human brain

The main goal of this work is to find an efficient way of reconstructing full genome regulatory networks and apply it to learning a GRN for the human brain. Full genome networks for humans can have from 20,000 to 50,000 nodes (depending on whether non-protein coding genes are considered or not). As discussed in the previous section, this size of network is usually very hard to work with due to the associated computational cost. Few algorithms scale well to this size, and most need HPC resources to be able to deal well with it.

As our dataset, we incorporated the microarray data from the Allen Brain Institute Human Brain Atlas. The dataset has measurements for 20,708 protein-coding genes with 3500 samples gathered from different areas of the brain.

#### Visualizing full genome networks

Our other main goal is to present a new tool for visualizing and interpreting massive BNs. The usual method used to visualize networks of this size is to decide sections of interest and show only subnetworks of a more reasonable size. Trying to visualize the whole network is usually unfeasible due to the computational expense required to render it and the way it can become exceedingly complicated to understand, thus losing the advantage of interpretability we are trying to achieve. To solve these problems, we implemented NeuroSuites-BNs (available at https://neurosuites.com), which we will explain in detail in the section *Visualization of Massive Gene Regulatory Networks*.

### FGES-Merge

FGES-Merge is our proposal for an algorithm capable of efficiently learning massive BNs without forcing initial structural restrictions. We broadly follow the structure of the algorithm from [43] but introduce several improvements that make our algorithm scale from the 1000-node networks tested in the reference algorithm to networks that are 20 times larger, such as the human genome network.

The structure of the algorithm is as follows: First, for each gene *X_i_*, we select its most likely neighbours as candidates for a local subgraph around *X_i_*. Next, we learn each local subgraph using a modified version of FGES. Finally, we merge the local subgraphs by performing graph unions with prunning to satisfy the topological properties of GRNs.

#### Neighbourhood Selection

We want to simplify the problem of learning a massive BN by dividing the network into a smaller neighbourhood network for each of the nodes and then merging them. To do this, we need to select which nodes will belong to each of the smaller networks. In [43], the authors calculate the pairwise MI between the nodes. Then, for each node *X_i_*, they assume the MIs come from two different distributions: one with the nodes in the neighbourhood of *X_i_*, and the other with the nodes that are not in the neighbourhood of *X_i_*. They assume that any MI sampled from the first distribution will always be higher than any MI from the second. This means that they can sort the MIs from highest to lowest and test the likelihood, for each possible size *s* of the network, that the first *s* nodes belong to a distribution and all the others belong to the second distribution and compare it with the likelihood that they are all sampled from just one distribution. Then, they take the most likely neighbourhood size *s_max_* and return the first *s_max_* nodes sorted by their MI with *X_i_* as the candidates for the first neighbourhood network. They then repeat this process for each of the nodes in the original graph (Fig. 2).

**Fig 2.**
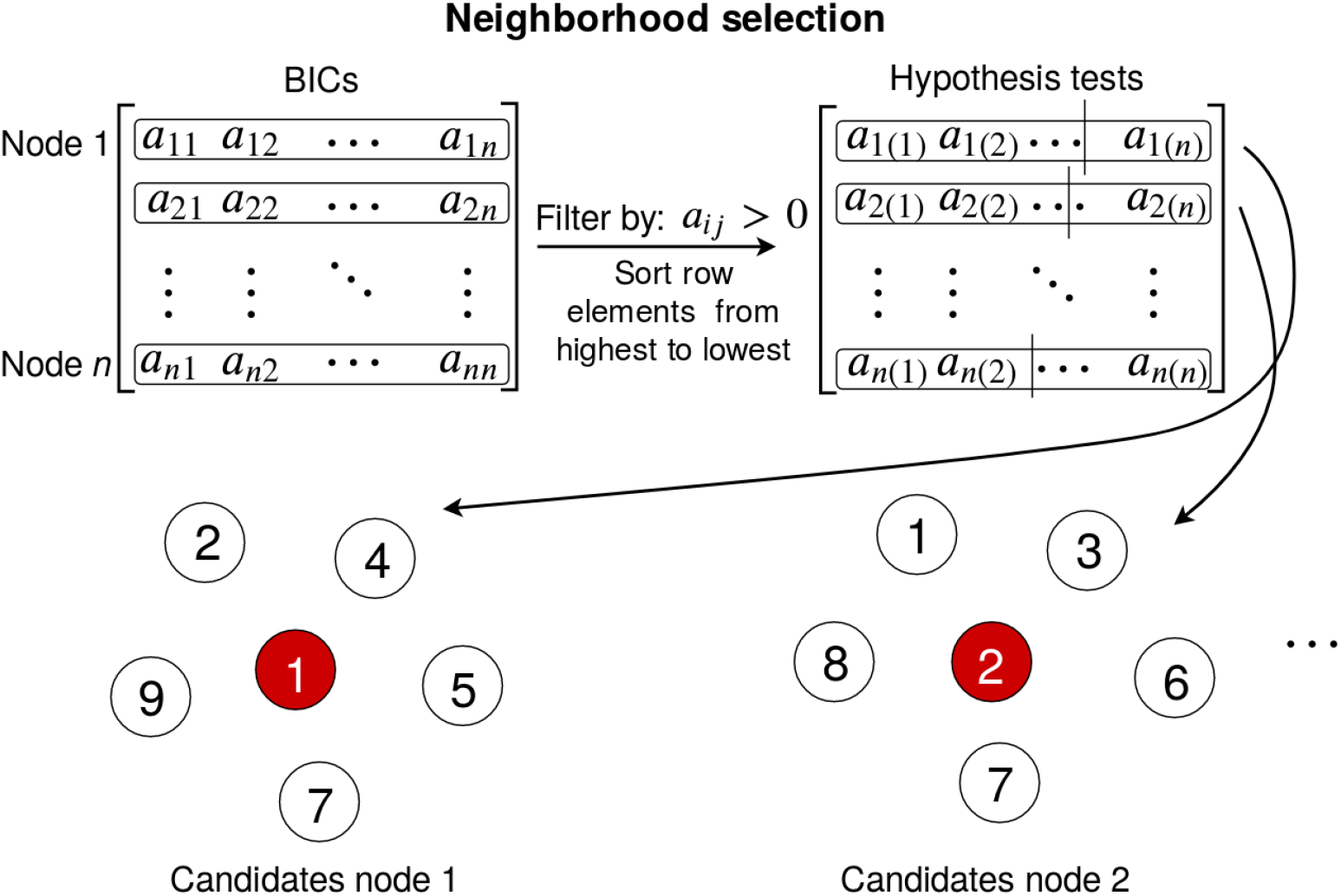
Diagram showing the neighbourhood selection step of the FGES-Merge algorithm. First the pairwise BIC matrix is computed by calculating the BIC score of adding the edge between each pair of nodes, such that every *a_ij_* corresponds to the BIC(*X_i_, X_j_*) (Eq. 7). Then, the BICs are filtered to only take into account the positive values and sorted from highest to lowest. We divide each row at the most likely point and take the left side as the neighbours for the next step.

We modify this procedure by changing our score from the MI of *X_i_* and *X_j_* to the local BIC [47] of adding the new edge from *X_j_* to *X_i_* (although this will be symmetrical at this stage), which is calculated with the following formula:

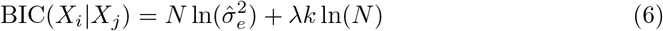

By using this expression for the BIC score, we assume that the level of expression of each gene is distributed as a linear Gaussian depending only on its parents, so 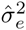 is the average of the sum of the squares of the residuals of regressing the child node against its parents. Here *n* is the number of instances, *k* is the number of parents and *λ* is the penalty which is a hyperparameter of the algorithm. As we saw in the review of the methods, assuming a Gaussian distribution for gene expression is valid and gives good results. It also speeds up the second step of the algorithm (FGES) because we use BIC again as a score for maximization, which is especially useful after we have found the structure since both estimating the parameters of the BN and performing inference become significantly faster.

We also do this because we will also need the BICs for the next step in the algorithm. Since calculating BICs is one of the most expensive steps and calculating MIs would take a similar amount of time without significant advantages, we replaced the values. of the BICs for the MIs in our version of the algorithm. Since the size of the BICs matrix is of order *G*^2^ and the calculations are independent of each other we parallelized this step.

Our second modification is limiting the possible size of the neighbourhood. We decided to do this because the topological properties of the GRN imply that the set of parents of each node is small. Even though the set of children can be very large for the hubs of the network, we assume that each child will contain the hub in its neighbourhood. This means that when we merge the edges between the hub and its children, we always expect the edge between the hub and each child to be added to the final network. This limitation makes each of the neighbourhood networks smaller thus speeding up the algorithm.

#### Fast Greedy Equivalence Search

In this step [43] uses a simple greedy strategy to learn the neighbourhood networks. Instead, we use our own variant of the FGES [12] algorithm which is much faster. FGES starts by greedily searching over the space of edge additions in the forward equivalence search (FES) step. At each step, it adds the best possible edge to the structure of the BN according to the BIC difference from adding that edge, which, if we want to add an edge from *Z* to *Y*, is given by:

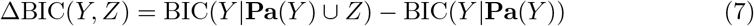

where each of the BICs is calculated using Eq. 6 after learning the multilinear regression of *Y* against the set of its parents (with and without *Z*). Then, instead of calculating the values of all possible edge additions again, our variant FGES algorithm uses the fact that since the BIC is a local score, the new edge can only modify the score of some of the edges around it. This allows us to skip many of the computations. Once no edge additions are possible (because they would all worsen the graph), we search the space of edge deletions in the backward equivalence search (BES) step to end up with the best scoring structure. (Fig. 3). Explaining the procedure in detail is beyond the scope of this paper, but the full pseudocode can be found in the appendix of [12]. The original algorithm is implemented in Tetrad [48], and our version can be found at https://gitlab.com/mmichiels/fges parallel production.

**Fig 3.**
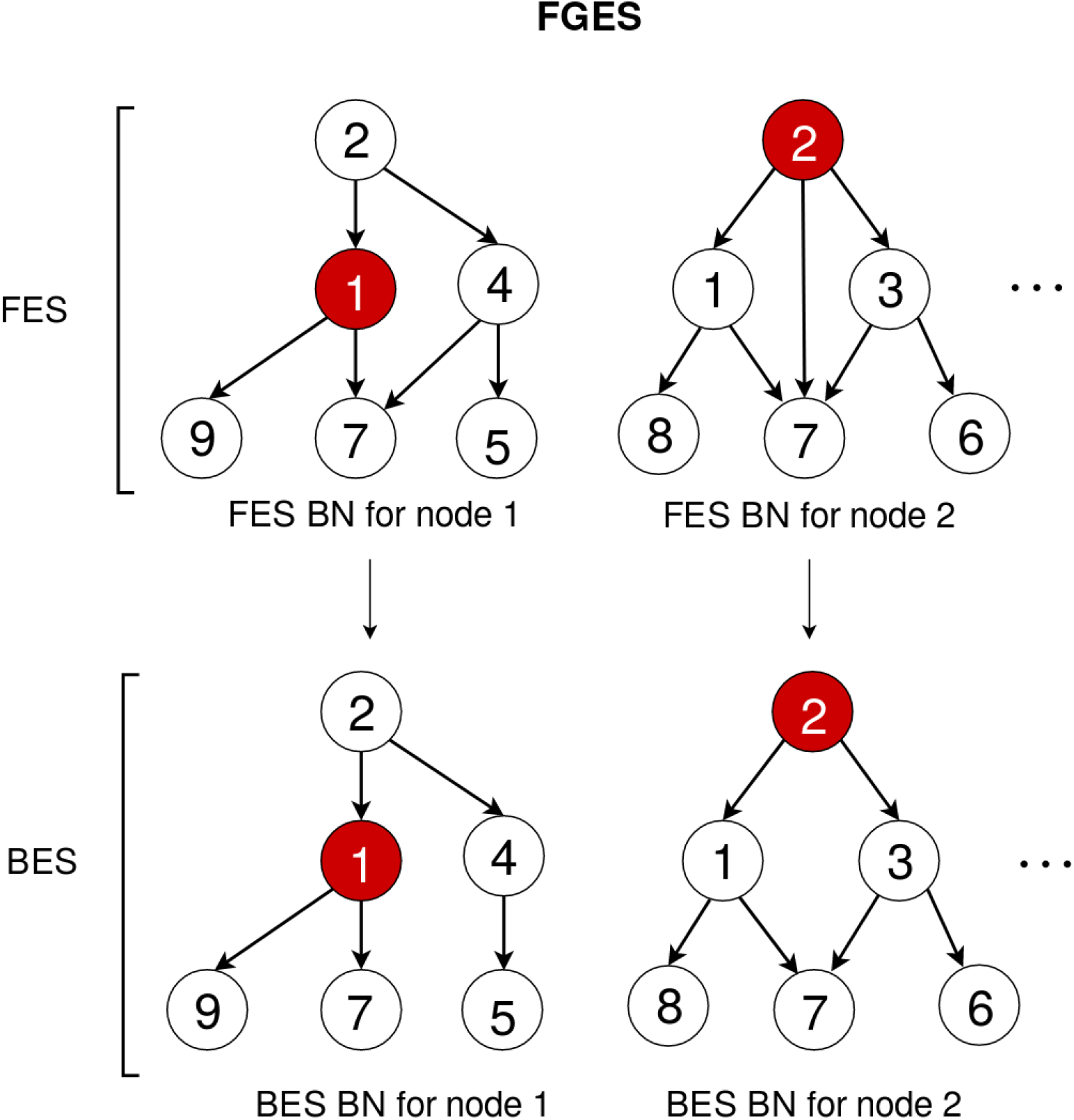
This diagram shows the basic flow of the FGES algorithm, following the example of Fig. 2. First, we take the neighbourhood candidates from the previous step and greedily (except for the simulated annealing step) add edges until the BIC cannot improve in the FES. Then, we perform any edge deletions that improve the score in the BES. This approach gives us the globally optimal neighbourhood network for the candidates. We then repeat the process for each of the other nodes.

The main modifications we do are parallelizing the calculation of the new possible edge additions at each step and adding simulated annealing to choose which edges to add or remove. The first change makes the slowest step of the original FGES algorithm much faster. We first create the set of all edges that have changed with the last addition and then divide it across all processors so that we can calculate their scores in parallel.

The second change sometimes chooses suboptimal additions and deletions to increase exploration by including a probability that a random addition or deletion will be chosen instead of the maximum scoring one. This probability will decrease with each iteration until it reaches zero, so the final steps of each network are always performed greedily. Given two neighbouring nodes X i and X j (in the sense of belonging to the same neighbourhood graph), we are usually going to find them together in many of the small networks. That is, if one of them is selected, the other is very likely to be selected too. Since the neighbourhood networks are smaller than the original, they are necessarily missing some context, so the optimal edges in the neighbourhood network might not be the same as those in the original. By allowing for suboptimal edges, we increase exploration in each of the subgraphs and make it more likely that the final structure contains the true edges. Since that structure will be pruned later, we do not have the problem of keeping edges that are very low scoring for the final graph.

#### Merge

The final step is combining all the learned local networks into a single global network for all the nodes. In [43], the authors try different methods for doing this and conclude that the best scoring method is to simply perform the union of all the graphs while checking for and removing any cycles. We found an improvement to this method by using a pruning strategy that removes the lowest scoring edges according to their BIC and the number of times they appear in the subgraphs. Since we only calculate the BICs once, at the beginning of the merging phase, this is an approximate strategy since we do not calculate the effect of each removal again. However, we found that the accuracy of the recovered network improved and the topology of the pruned network more closely resembled the true topology of a GRN.

We also added a step that allows the user to optionally introduce a predefined list of hubs (which in the case of GRNs would usually correspond to transcription factors) so that edge orientation can be consistent with expert information. If this list is missing the algorithm chooses the nodes with the highest number of neighbours as hubs and, if we find inconsistent orientations during merging, makes the hubs the parents. Again, we found that this change improved the accuracy of edge orientation and made the topological properties of the BN more closely match those of a true GRN. See Fig. 4 which summarizes the merging and pruning process.

**Fig 4.**
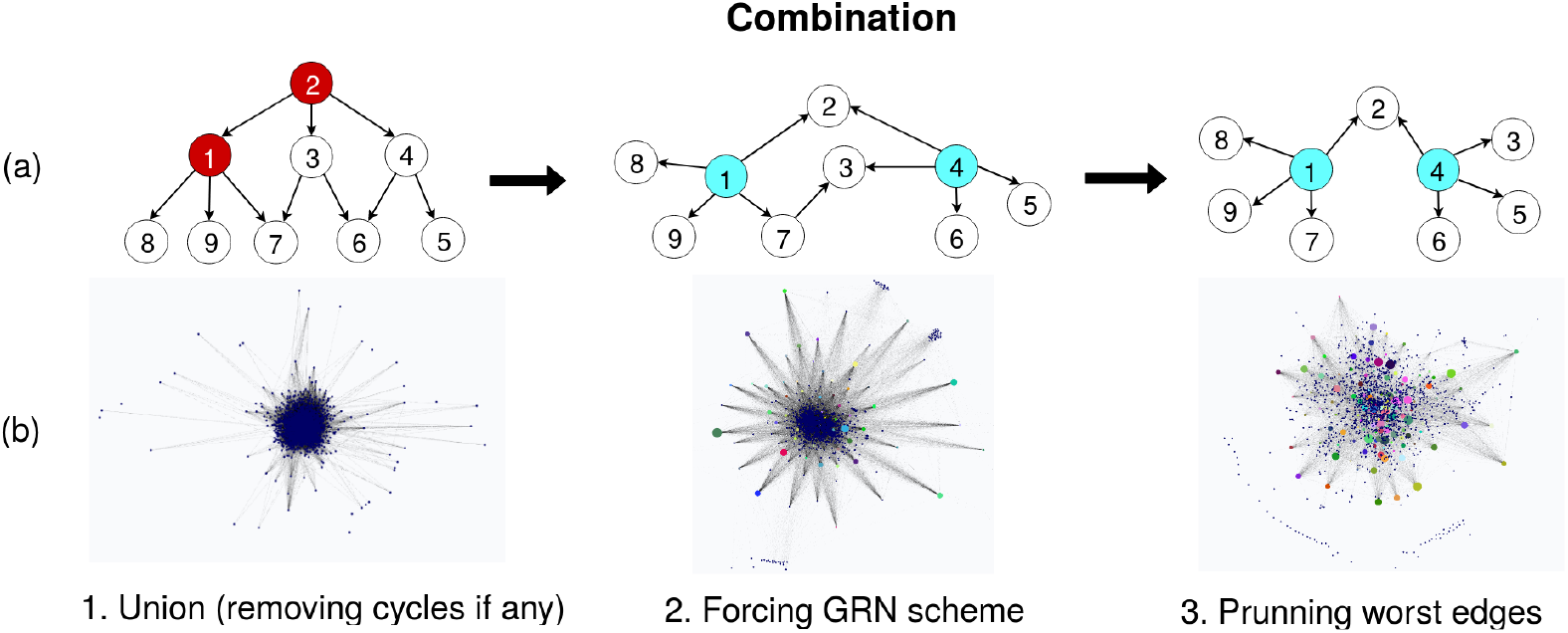
Diagram explaining the combination steps. (a) Simplified example continuing from Fig. 3. First, we merge the neighbourhood networks with their union, removing any existing cycles. In the second step, we find the hubs (in blue) either via expert knowledge or by using the most-connected nodes in the network to orient the edges from the hubs to their connected nodes that are not also hubs while checking for cycles and removing them. Finally, in the third step we prune the worst performing edges by their BIC score up to a size threshold corresponding to the expected number of edges in a GRN (approximately # edges *<* 10*G*, see [7]). (b) Real world use case of the combination procedure with our learned human brain network.

### Parameter Learning

The CPD of a node *Y* with parents **Pa**(*Y*) in a Gaussian Bayesian network is given by:

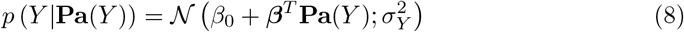

To estimate the parameters *β*_0_*, **β***, and 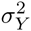 for each node, we learn a multilinear regression against the set of the parents of the node. The parameters of the regression give an estimate of the coefficients of the mean of the CPD for the node and the mean of the sum of squares of the residuals of the regression gives an estimate of the variance.

### Inference

Exact inference in a Gausian Bayesian network is easily done since these networks represent multivariate Gaussian distributions. If *Y* is a linear Gaussian of its parents (i.e., it follows Eq. 8) and **Pa**(*Y*) are jointly Gaussian with distribution 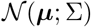, then the distribution of *Y* is a normal distribution 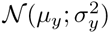 with

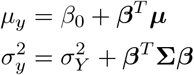

And the joint distribution over **Pa**(*Y*) and *Y* is a normal distribution where:

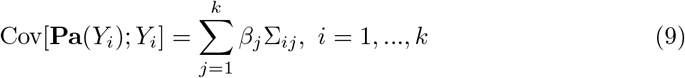

We can convert from one representation to another by ordering all the variables topologically (no child is before their parents) and iterating over the topological order, adding nodes one by one to the joint distribution.

Then, we have two operations: marginalization, which gives us the joint distribution of some variables of interest and conditioning, which sets the values of some of the variables and returns the conditional distribution given those values.

For marginalization, we just need to extract the means, variances and covariances of the variables of interest. For conditioning, we follow the procedure described by Koller and Friedman in [35] which requires converting the Gaussian into information form (for which we need to invert the covariance matrix) and setting the values of the evidence variables. We do not need to invert the whole matrix, just the block containing the evidence variables. To obtain the posterior CPD for the variable *Y* after conditioning on a set of variables ***X*** we use the following equationsa:

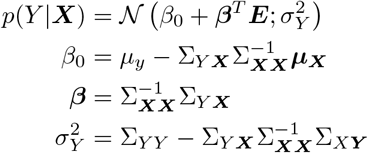

In our implementation we vectorize the formulas to obtain the parameters for the |*G*| − |*X*| variables left after conditioning with just a single matrix multiplication. Normally, inference in BNs is exponential in the number of nodes, but for Gaussian Bayesian networks, it is reduced to matrix multiplication and inversion so the complexity is *O*(*l*^3^) where *l* = max(|*E*|, |G| − |*E*|), which makes the complexity tractable for our 20,000 node network.

### Visualization of massive gene regulatory networks

Genome-wide GRNs are on the order of thousands of nodes and edges and hence are difficult to visualize. However, as for every other network, visualization is a key aspect for understanding and analysis. There are two main concerns that need to be overcome to develop a visualization tool for massive networks: computational efficiency and usability. One common method to alleviate the computational burden and make the network sufficiently tractable in order to use common tools for the analysis of BNs is to show only subgraphs instead of the global network. While this approach is valid for showing some relevant genes and their connections, we would prefer to have the ability to visualize and work with the whole network at once.

We used the Sigma library for the graph visualization task. Sigma is a JavaScript library for graph visualization that provides a WebGL backend to make use of the GPU resources. GPUs have substantially increased in power in recent years and are now able to solve the problem of visualizing a massive network. Since the library is a general-purpose graph visualization package, we implemented all the necessary modifications to adapt it for BNs. That is, for proper visualization of BNs, we need to visualize the node parameters, run inference-related operations such as making queries and observing the posterior distributions, implement specific highlighting tools such as showing the Markov blanket of a node, etc. In summary, we need a rich set of interactive tools to fully understand the BN structure and parameters. This is where current BN visualization software frameworks fail for massive BNs, as their implementations of these operations are not scalable to tens of thousands of nodes.

The goal was to make a complete modern solution for BN learning and visualization, so we developed a web application to include all the desired functionalities. The main advantages of this approach are the ease of use, the interactive capabilities and computational efficiency. The ease of use is achieved by the software architecture, as a web application is inherently easier to access than desktop software where the user may need to install multiple packages or different dependencies for every operating system or hardware architecture. Furthermore, the user interface has been specially designed to manage massive networks, resulting in well-polished interactive tools for working with the BN, as we explain below. The whole web service was designed with computational efficiency in mind, having been optimized for the visualization task from the beginning, including a separation of the visualization code from the business logic code for managing the graph algorithms (such as computing different layouts).

#### Interactive tools for BN visualization

Now, we will focus on the interactive tools we developed to be able to understand massive BNs. Some of these tools are general purpose graph tools, while others are specific for BNs. One of the most important general purpose tools is the selection of layouts to position the nodes and edges in a meaningful way. It is possible to run every layout for the BNs, but force-directed layouts such as the Fruchterman-Reingold or ForceAtlas2 algorithms [49] are recommended for GRNs, as they enable us to notice the formation of some clusters (see Fig. 5 to view a selection of the available layouts).

**Fig 5.**
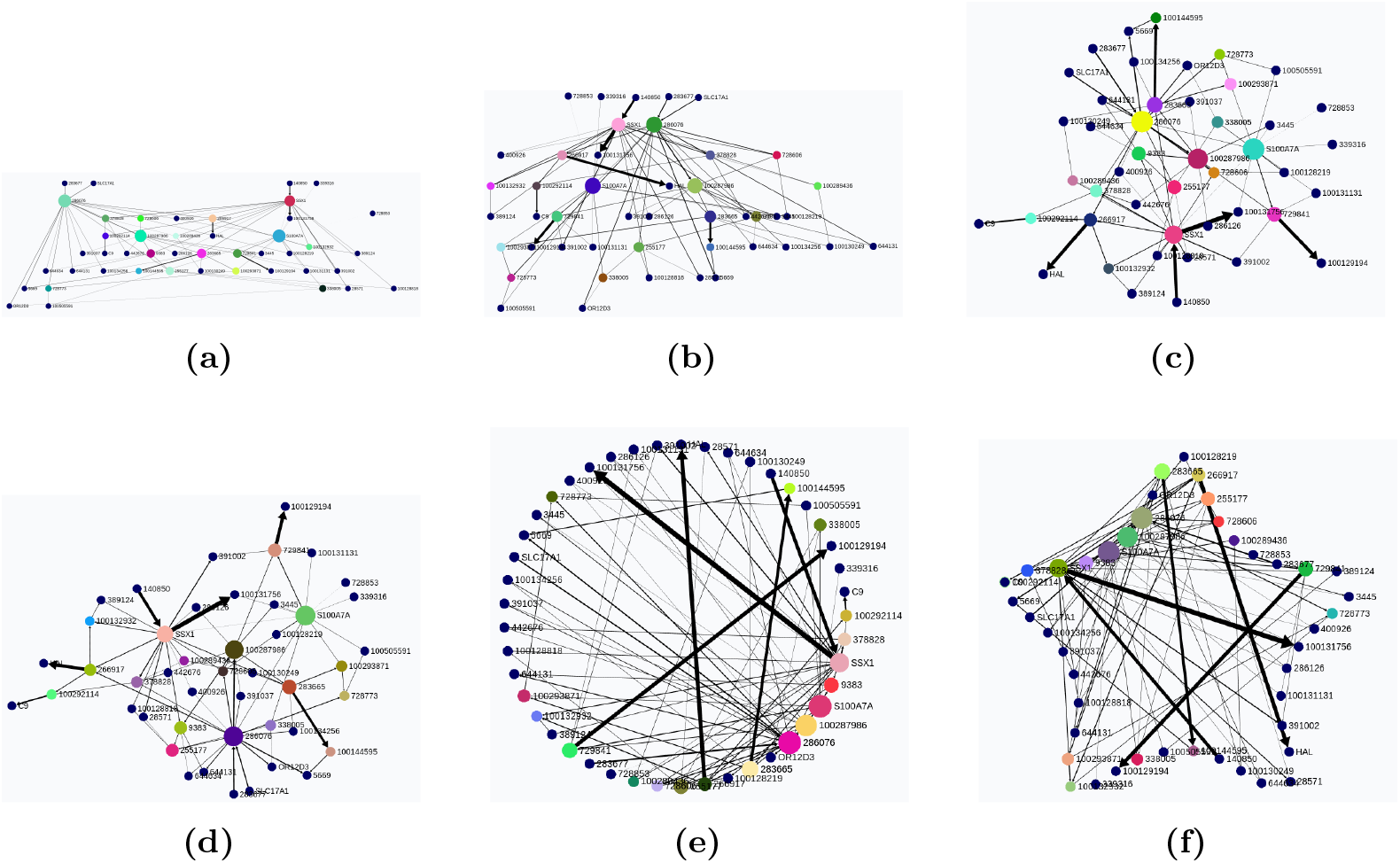
Some layouts algorithms for BNs visualization, highlighting the important nodes with the betweenness centrality algorithm. (a) Dot layout. (b) Sugiyama layout. (c) Fruchterman-Reingold layout. (d) ForceAtlas2 layout. (e) Circular layout. (f) Image layout of a star shaped picture.

To understand massive networks, we usually need multiple ways to find and select the desired nodes. To highlight important nodes there are two main options: a user defined list of nodes ordered by groups or a set of automatic detection algorithms. These algorithms can either highlight nodes by some topological properties, such as their degree or betweenness centrality, or group them by communities by running the Louvain algorithm [50]. Usually, the combination of a force-directed layout with the Louvain algorithm provides interesting insights into possible clusters in the network structure.

To provide a use case for dealing with a user-defined list of groups, we downloaded the DisGeNet genes-diseases metadata database of [51]. his provides information for grouping the genes by diseases they are associated with, making it useful for incorporation into the visualization of our learned GRNs. The user can select a specific disease and view all the associated genes. Conversely, the user can select a specific gene and view the disease associated with that gene as well as all the genes associated with that same disease (Figs. 6a and 6b).

**Fig 6.**
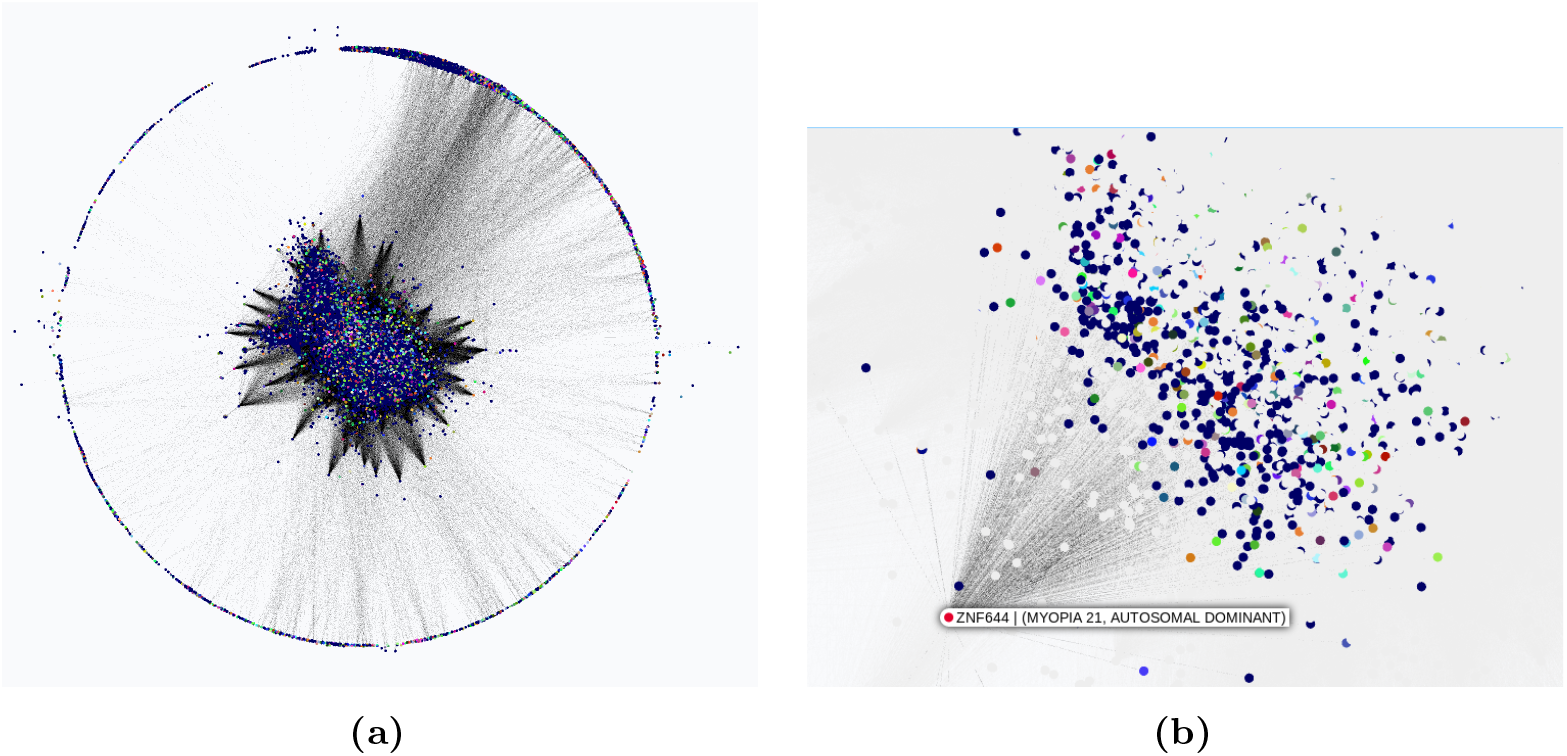
Nodes colored by the genes-diseases metadata association from the DisGeNet database. Our learned full brain human network. (a) All the genes colored by their main associated disease. (b) Node ZNF644 selected to view its disease association (Myopia) and all its children genes related to it.

The tools we developed specifically for BNs are for visualizing parameters and performing probabilistic inference. In our Gaussian Bayesian networks, the parameters are shown as an interactive plot of the marginal Gaussian distribution, where the user can zoom and hover the mouse over the plot to see the distribution values.

To perform inference, the user must first set a specific value for a node or a group of nodes. This will set the selected nodes as evidence variables ***E*** and give them a special colour (red).The evidence variables, ***E***, can be selected in three ways: selecting one specific node as evidence (Fig. 7a), selecting a group of nodes as evidence (from an imported user defined groups file or Louvain algorithm) and defining a new list of nodes.

**Fig 7.**
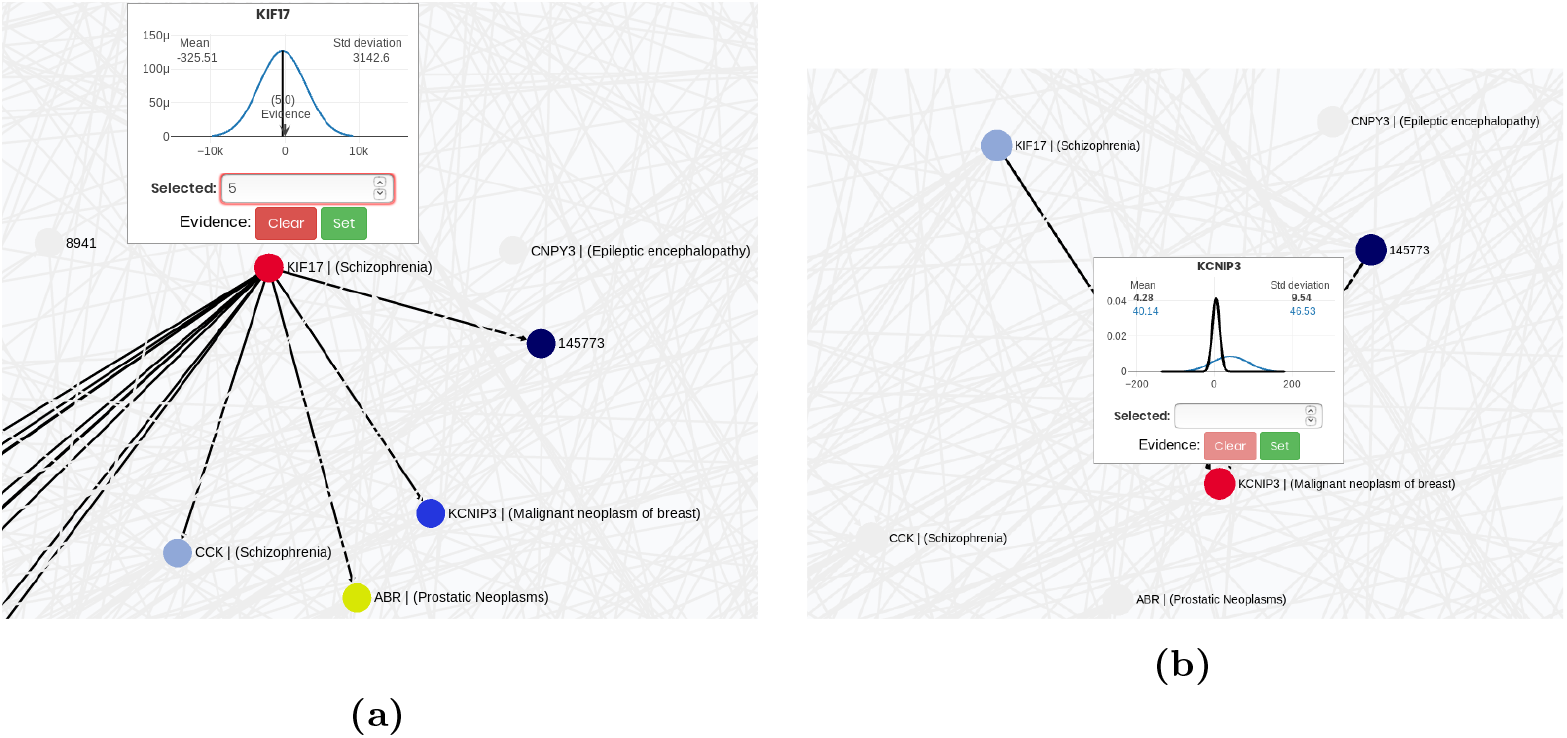
Inference workflow in the BNs visualization. Our learned full human brain network. (a) Set the evidence *E* = *e* in a specific node, in this case the one in red, corresponding to the KIF17 gene associated with the schizophrenia disease. (b): *p*(*Q E* = *e*), the posterior distribution of a node *Q* (the one at the bottom in red, corresponding to the gene KCNIP3 associated with malignant neoplasm of breast) given the evidence, where *e* is the KIF17 gene. In this case the network have found a relationship between these two diseases, since a very low value near zero for the gene associated with schizophrenia (KIF17) results in a very low value for the gene associated to malignant breast neoplasm (KCNIP3). We have found some evidence for this in the literature [52].

The inference is performed on the server side to provide an efficient implementation; therefore, we do not need the user to have a powerful computer that could help with usability. This allows us to run the inference in less than 30 seconds, even in a massive network with 20,000 nodes. The resulting multivariate joint distribution and the original joint distribution are cached in the back end to provide a faster visualization of the results. This means that the process is almost invisible to the user and is done only when evidence or a group of evidence is fixed.

The second step is to visualize how the distributions of other nodes have changed with respect to the original ones before setting the evidence values. The user can either click on or search for a specific node or select a group of query nodes ***Q*** to obtain the posterior *p*(*Q E*). his will be shown as an interactive Gaussian probability density function plot in blue, while the original distribution will be shown in the same plot in black (Fig. 7b).

Following the previous gene-disease use case, we can set as evidence nodes all the genes associated with a specific disease. In this way, we can, for example, set a high value for these genes and see how the distribution of genes associated with another disease has changed.

Finally, to support visual differential analysis, we created a tool that allows for comparison between two networks. This tool works by overlapping two networks with the same nodes so that the differences between the edges are shown with different colours, as in Fig. 8. We also provide a tool for showing only the edges of the first network, the edges of the second network or the common edges.

**Fig 8.**
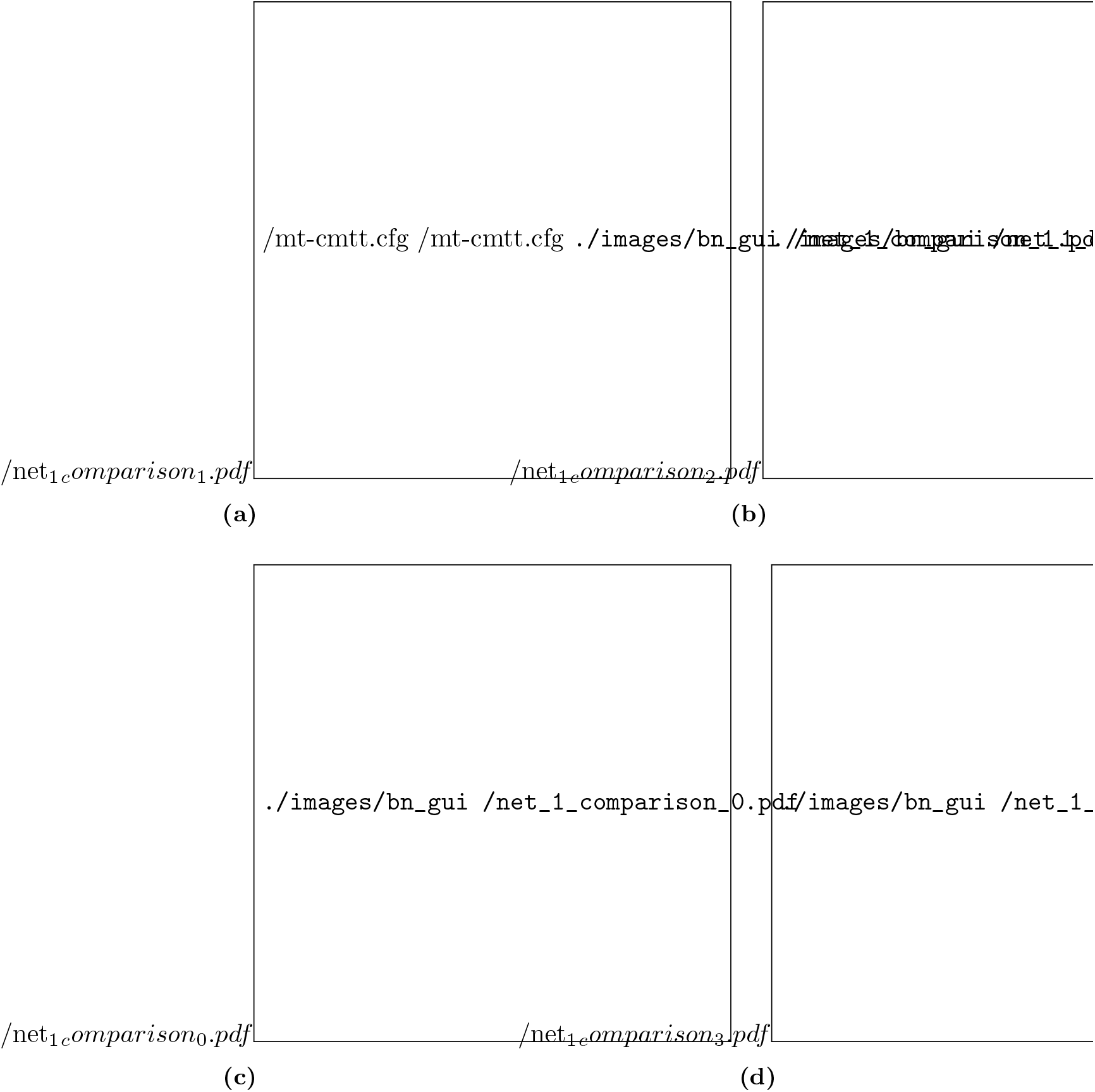
Structure comparison of the Network 1 of the DREAM5 challenge. (a) True graph edges. (b) FGES-Merge learned graph edges. (c) All edges. True graph edges are displayed in black, FGES-Merge learned graph edges in blue and common edges in green. (d) Common edges between the True graph and our learned graph with FGES-Merge.

All the software has been packaged as a Docker [53] container to provide a production ready solution. Since the computationally costly code is running in our backend, the users only need an average GPU card to learn and visualize massive BNs fluently.

## Results

### Benchmark tests

We wanted to compare the ability of our method to recover the underlying structure of GRNs against that of a variety of algorithms, even those that do not use BNs. However, we also wanted to compare our method to other BN methods as a way to show that our algorithm transcends the usual difficulty in learning BNs and is capable of scaling to thousands of nodes.

To achieve these comparisons, we ran two different tests. First, we compared our accuracy in recovering the structure of the networks from the DREAM5 challenge [28]. We used networks 1, 3, and 4 following [28], since network 2 was discarded during the original challenge. The second test was against other common methods of learning BNs with the same data, but instead of measuring accuracy, we measured the time taken to learn the structure.

All the experiments for both the benchmarks and final results were run in our MPI cluster with three nodes, each one running in Ubuntu 16.04, Intel i7-7700K CPU 4 cores at 4.2 GHz, and 64GB RAM.

#### Structure recovery benchmark

We used the DREAM5 challenge results repository avialble at https://www.synapse.org/!Synapse:syn2787211 to obtain the structure learned by all the original competitors. This allowed us to compare our results with those of the competitors without having to run all the algorithms ourselves, which might have been impossible since not all of the other methods are available and we expect that some of them would have taken a very long time to run.

Each of the methods for the benchmark outputs a matrix of G × G entries, one for each possible edge in the network. Each entry corresponds to the probability assigned to the edge by the method. BN methods do not usually do this, and we believe that this might have been one of the reasons they performed so poorly during the original challenge. Our solution was to establish a threshold for all the methods and transform their probabilities into a series of binary predictions. To do this, we had to take into account that the network is sparse, so the prior probability for the binary classification problem is not 0.5 but instead approximately 1/number of nodes (since we expect the number of edges and nodes to be the same order of magnitude). Had we not done this, any method that was well calibrated and used the proper prior would get close to 99

The original score for the DREAM5 challenge was the area under the precision-recall curve (AUPRC), which is usually a good score for imbalanced problems. However, it presented two problems. First, it was hard for us to implement more than one threshold since, as stated above, BN methods do not give probabilities to edges, so we could not calculate the AUPRC properly for our method. Second, since the method uses multiple thresholds, a good score here does not directly translate to its usefulness in the lab.

When given one of these networks, a biologist has to make a choice on what interactions to test along with the corresponding cost, and they will not have the luxury of knowing which of the thresholds will give the best results for the network beforehand. AUPRC is still used because it maximizes a combination of precision and recall and deals well with class imbalance problems. In the end, we decided to use a score that has these two properties and does not have the problem of using many thresholds, the Matthews Correlation Coefficient (MCC) [54], which is basically an extension of the F-score to deal with class imbalance. The expression for the MCC from the confusion matrix is:

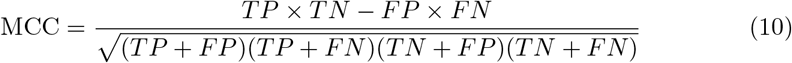

where TP, TN, FP, and FN are true positives, true negatives, false positives and false negatives respectively. The MCC ranges from 0 to 1 with 0 meaning that the classifier has misclassified a whole class (predicting all edges or all non-edges) and 1 meaning perfect classification. Fig. 9 shows the MCC scores for all the methods in the DREAM5 challenge and FGES-Merge for networks 1, 3 and 4. FGES-Merge does better than every BN method except 1 and 2 (simulated annealing with Catnet in R) as seen in Fig. 9a (in which FGES-Merge is one of the best performers overall) and Fig. 9b, (see Fig. 8d to see how much of the actual structure was recovered). Furthermore, in Fig. 9c, where BN methods 1 and 2 have very poor scores, FGES-Merge is still one of the best methods. Fig. 9c highlights the most complicated task of the three since the task involves the GRN for *Saccharomyces cerevisiae*, a eukaryote, which has more complex gene regulation processes compared to *E. coli*, a prokaryote, in network 3, or the *in silico* network in (a) which is also modelled after *E. coli*. We expect this good performance for eukaryotic cells to translate well to the human brain GRN.

**Fig 9.**
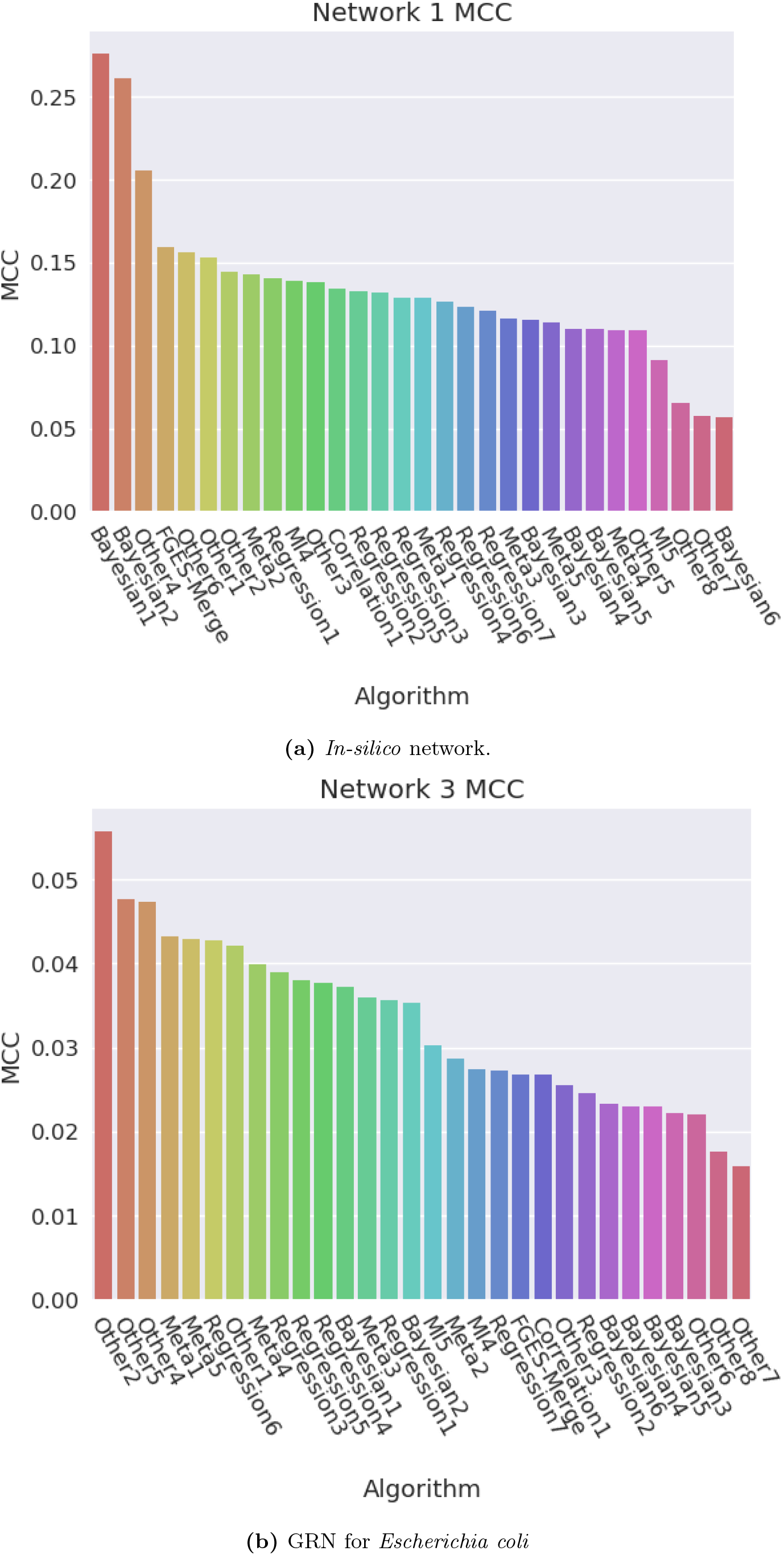
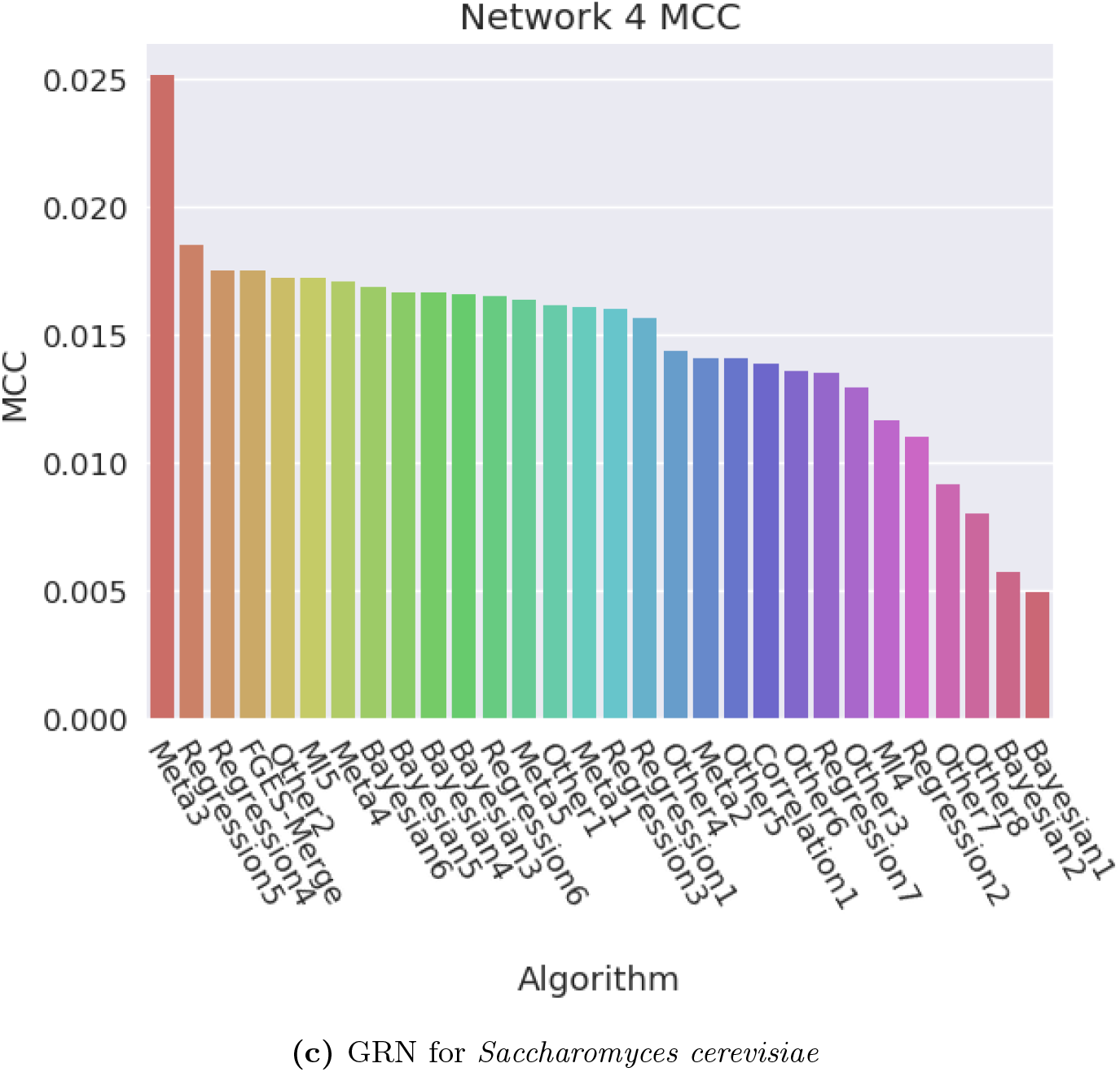
MCC scores for all the methods in the DREAM5 challenge and FGES-Merge for networks 1, 3 and 4. FGES-Merge does better than every BN method except 1 and 2 (the simulated annealing with Catnet in R) in (a) and (b), being one of the best methods in (a). Furthermore, in (c) where methods 1 and 2 of BNs have very poor scores, FGES-Merge is still one of the best.

#### Times benchmark

For our times comparison, we tested some of the most common BN learning algorithms implemented in the bnlearn R package [55] and one regression method (GENIE3) on the smallest network (≈ 1000 nodes) from the DREAM challenge (network 1). We attempted to show whether our algorithm scaled better than the others by running all of them in increasingly larger networks, expecting that our algorithm would be beaten by Chow-Liu’s algorithm since it has a complexity of *O*(*G*^2^) and probably by GENIE3, since we expected regression to be faster than learning a BN. In the end, we could not test more than the smallest network since although our method finished in slightly more than an hour, the other non-quadratic BN methods ran for more than a day and did not finish.

The results present in Fig. 10 show that our algorithm is slower than that of Chow-Liu, as expected, but slightly faster than GENIE3 which we did not expect. These results are surprising because they imply a massive improvement in speed between FGES-Merge and most other BN learning algorithms with very few restrictions.

**Fig 10.**
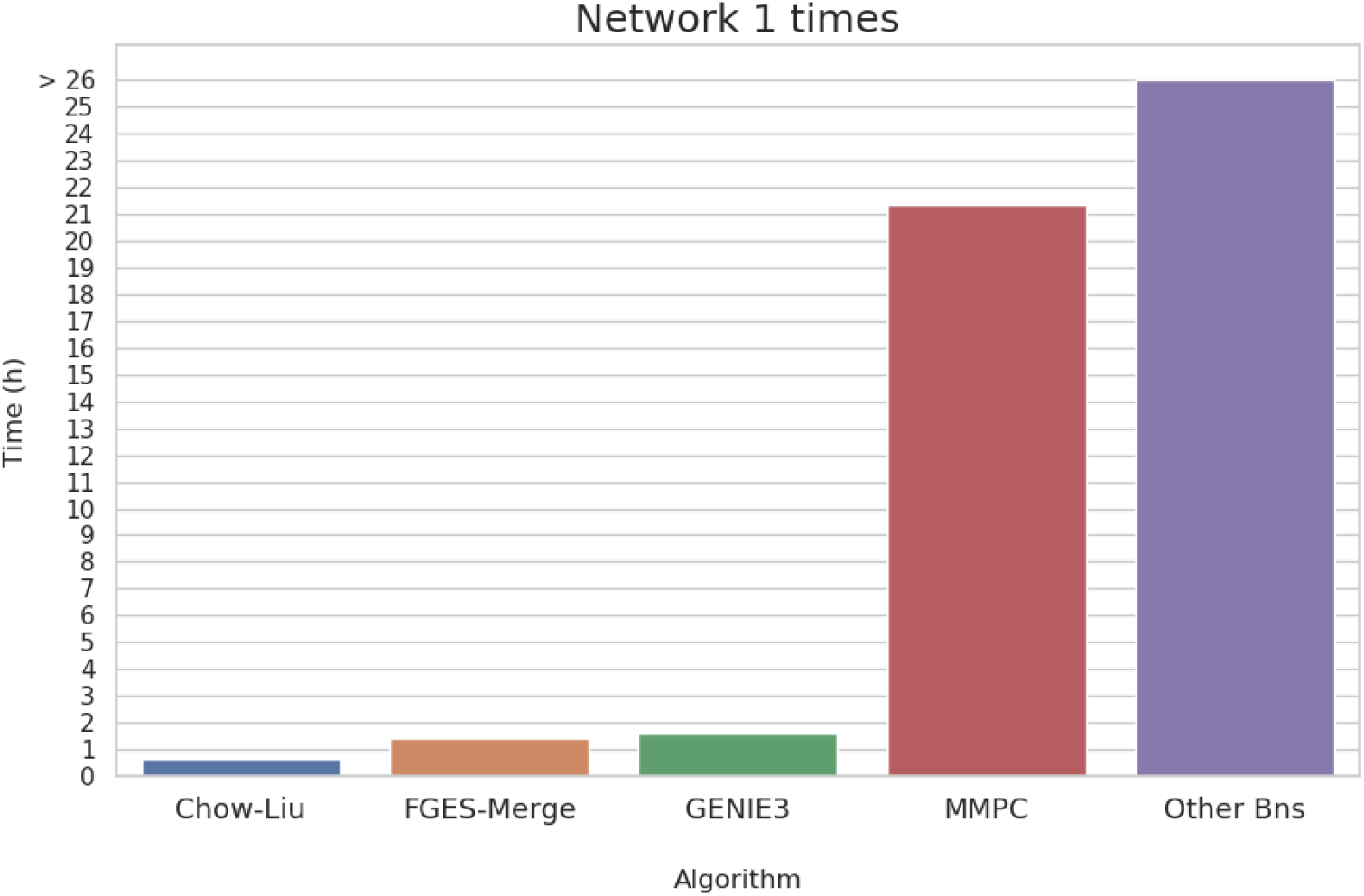
Time taken (in hours) for different methods of learning a GRN for network 1 of the DREAM5 challenge. Among the BN learning methods only Chow-Liu’s algorithm is competitive in time which is normal since its complexity is *O*(*G*^2^). Even a regression based method like GENIE3, which is supposed to be much faster than BN based methods, is slower than FGES-Merge. Algorithms that did not finish before the 26 hour mark were forced to stop early.

### Human Brain Regulatory Network

Finally, we applied our method to solve the problem we were first interested in: learning the genome-wide regulatory network for the human brain.

The main objective was to find genome-wide GRNs for various areas of the brain (the cortex, white matter, cerebellum, hypothalamus, etc.). These networks would be very useful tools for biologists who are interested in studying how the functional differences between brain areas might arise from differences in genetic expression. To achieve this, we started with the Allen Human Brain Atlas microarray data and filtered only the protein coding genes. Then, we used FGES-Merge to obtain a series of GRNs for some of the high-level structures defined by the ontology of the Human Brain Atlas. We decided to only obtain the GRNs of these high-level structures instead of performing a more fine-grained approach because we did not have enough samples in our dataset to guarantee good results for smaller structures.

The application of FGES-Merge allowed us to achieve all of our objectives. We were able to obtain networks for various areas of the brain and an extra network for the average expression of the whole brain. In this section, we will show some of the generated networks, comment on possible applications of our models and discuss the topological properties of the learned networks to see if they respect the known empirical properties of GRNs.

#### Networks learned

Fig. 11 shows two networks obtained with the whole dataset, that is GRNs for the average gene expression level of the whole brain. These GRNs were learned with different penalty parameters for FGES-Merge (see Eq. 6) and thresholds for the number of edges. We can readily see how the higher-penalty network (Fig. 11a) presents various disconnected components unlike the lower-penalty network (Fig. 11b). This is what we would expect since a higher penalty forces sparsity. The second network predicted more edges than expected so the pruning step of FGES-Merge was used to remove some of the worst edges.

**Fig 11.**
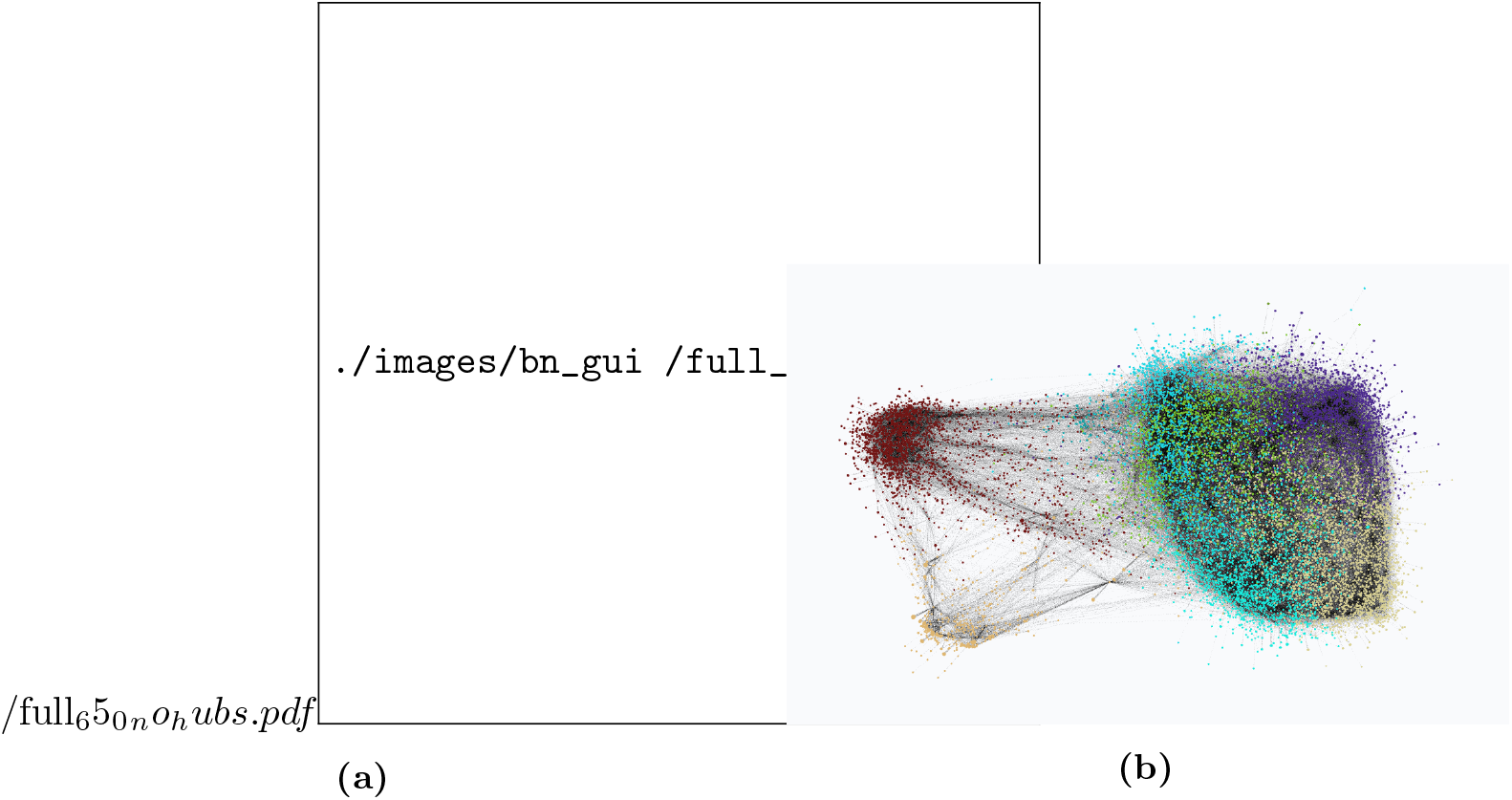
Human brain genome networks. The figure shows two networks learned with the whole Allen Brain Atlas dataset with different parameters for FGES-Merge. Both are visualized using NeuroSuites-BNs and colored via the Louvain algorithm to identify groups pf genes. (a) Network learned with a high penalty (*λ* = 65) for the FGES-Merge algorithm. (b) Network learned with a lower penalty (*λ* = 45) for the FGES-Merge algorithm

These networks can be used to visually search for relationships between genes, such as those shown in Fig. 6, which can then be tested to obtain possibly useful information about diseases or development. It can also be used quantitatively as in Fig. 7 to obtain concrete predictions about the effects of some genes on other genes’ expression levels. This could complement clinical trials that aim to alter gene expression by helping researchers decide which genes might be good targets for medication and to explore possible side-effects.

Finally, multiple networks for different areas could be visualized as in Fig. 8 to serve as an aid for differential analysis. Instead of observing only the gene expression levels between different conditions, we could see the structure and parameter differences, obtaining a clearer picture of the differences in gene regulation that lead to differences in gene expression.

### Topological properties of the learned GRNs

Fig. 12 shows the in-degree and out-degree distributions for the human brain GRN and the tail of the total degree distribution (the approximately 1000 nodes with the highest degree). As we discussed before, we expect most nodes to have an out-degree of 0 (they are not regulators), but the nodes that have children should have many of them, as expected from the evolutionary argument. Fig. 12b shows exactly this: of the 20700 nodes, most have zero children and so do not appear in the histogram. The ones that do have children can have over 1500 with an average of over 600 children.

**Fig 12.**
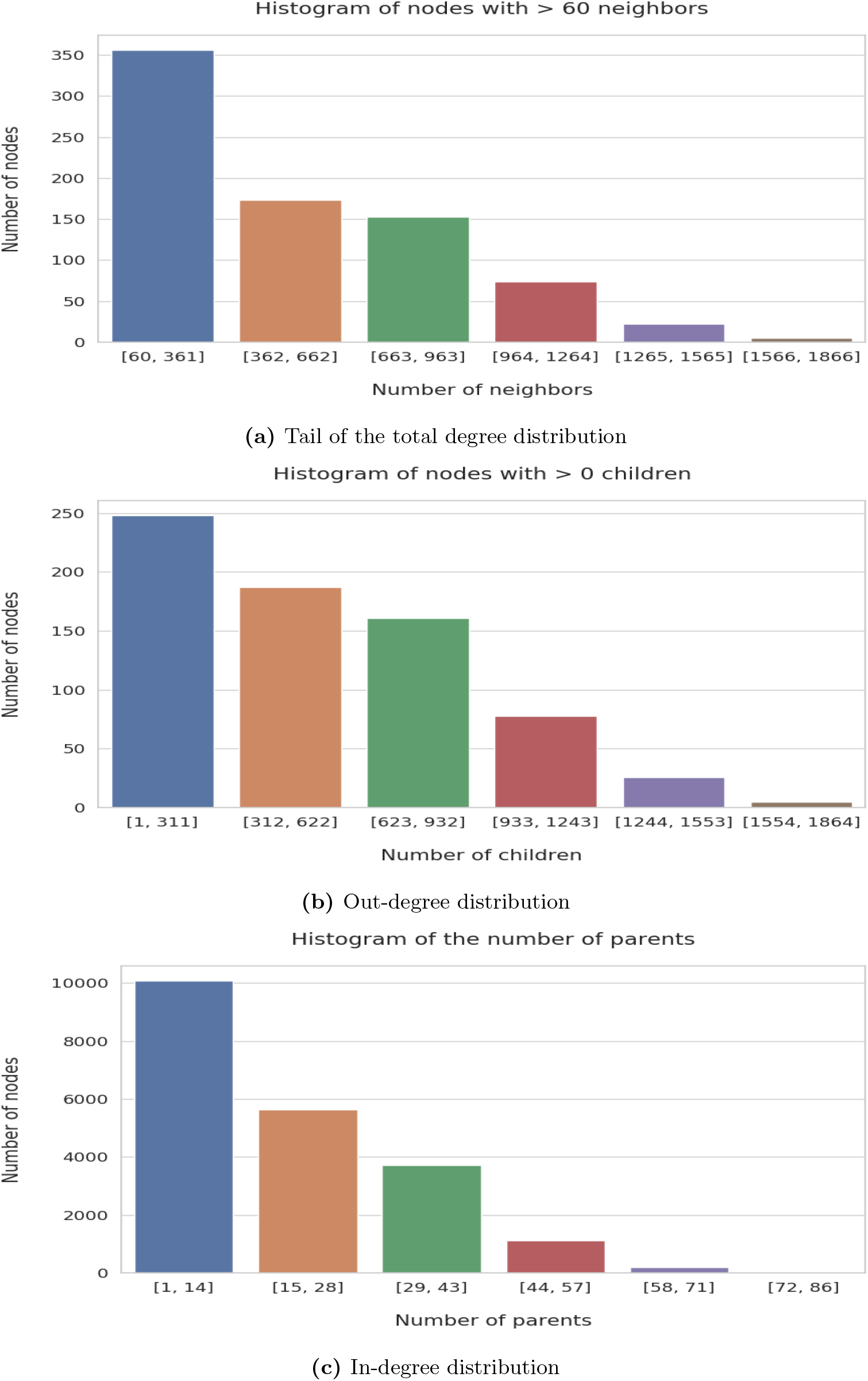
Node degree histograms for our learned full human brain network. The threshold for the minimum number of neighbors and children has been adjusted for a proper visualization of the histograms. The network corresponds to the Fig. 11b.

The in-degree distribution in Fig. 12c is also as expected. Most genes are regulated by a small number of regulators, and even the most regulated do not have more than 80 parents. Again, this is in agreement with the biological argument that gene regulation requires physical interaction, indicating that there is not enough room for hundreds or more regulators.

Finally, from all three distributions, we can see that the network is globally sparse but locally dense. Most nodes have no children, as seen in (b), and approximately half of the nodes have fewer than 14 parents, as seen in (a), with most having just one or two. However, the tail of the total degree distribution in Fig. 12a shows that the 1000 nodes with the highest degree have an average of 500 neighbours, with some of them having well over a thousand. That is, even though most nodes have almost no neighbours and the total number of edges is small, there are nodes that have very dense neighbourhoods. These results show that our algorithm respects the topological properties of GRNs, which gives us some reason to believe that the inferred network structure is sound.

These results show that our algorithm respects the topological properties of GRNs. This gives us some reason to believe that the inferred network structure is sound.

## Discussion

### Conclusion

In this work, we have reviewed the problem of reconstructing GRNs from gene expression data. In general, the problem is very difficult, and even the best methods have low scores on the reconstruction. Furthermore, every method that tries to scale to genome-wide networks while creating a quantitative predictive model will have to use HPC or make strong assumptions on the structure and parameters of the inferred GRN.

We opted somewhat for this second option, restricting the expression levels to be linear Gaussian distributions to be able to learn a BN, which would have the advantage of being very easy to interpret for experts who might want to use our tool, and we were able to scale to genome-wide networks by parallelizing the most time-consuming parts of the algorithm. The method we have presented, FGES-Merge, is competitive with the state of the art while also beating most BN methods and giving consistently good results even for the harder networks in the benchmarks. Our method is also much faster than any competing BN-learning methods. Furthermore, FGES-Merge gives results that respect the topological properties of real GRNs.

Our choice of model also has the advantage of immensely reducing the time required to perform probabilistic inference in a network of this size, making it much more useful for any kind of biological research since it is able to answer queries almost immediately. Finally, we also presented a new software tool that incorporates the learning algorithm FGES-Merge, the inference algorithms and the visualization tool NeuroSuites-BNs, which is available at https://neurosuites.com.

The source code for the FGES-Merge algorithm can be found in our repository: https://gitlab.com/mmichiels/fges parallel production. The code for NeuroSuites-BNs can also be found in our repository: https://gitlab.com/mmichiels/neurosuite. All our obtained networks learned from the DREAM5 challenge and from the full human brain genome can be found in the following folder in our repository: https://gitlab.com/mmichiels/fges parallel production/tree/master/BNs results paper

## Acknowledgements

The authors would like to thank Sergio Paniego for his help in the development of the NeuroSuites-BNs visualization tool.

This project has received funding from the European Union’s Horizon 2020 Framework Programme for Research and Innovation under Specific Grant Agreement No. 785907 (HBP SGA2).

## Notes

https://gitlab.com/mmichiels/fges_parallel_production/tree/master/BNs_results_paper

### Competing Interest Statement

The authors have declared no competing interest.

